# No evidence for intermolecular recombination in human fibroblast and blood mtDNA from individuals with biparental mtDNA transmission

**DOI:** 10.1101/2020.02.26.941922

**Authors:** Baoheng Gui, Zeyu Yang, Shiyu Luo, Jesse Slone, Sushma Nagaraj, Lisa Sadzewicz, Luke J. Tallon, Taosheng Huang

## Abstract

Strictly maternal inheritance and lack of intermolecular recombination of the human mitochondrial genome (mtDNA) are the assumed preconditions for molecular evolution studies, phylogenetic reconstruction and population genetic analyses. This hypothesis, however, has been challenged by investigations providing evidence for genetic recombination of mtDNA, thus sparking controversy. Using single-molecule real-time (SMRT) sequencing technology, we sequenced the entire mtDNA from blood and fibroblast cells from five individuals with biparental mtDNA transmission in three separate, multiple-generation families. After phasing the single nucleotide polymorphism (SNP) genotypes of mtDNA, no intermolecular recombination between paternal and maternal mtDNA was found when the mtDNA was transmitted in either biparental or maternal mode. Our study provides support for the argument that intermolecular mtDNA recombination is absent or extremely rare in humans. As a consequence, these results support the feasibility of mtDNA-based molecular evolution studies and phylogenetic and population genetic analyses for humans, while also avoiding inaccurate phylogenetic inferences and incorrect rejection of the molecular clock.

---

The human mitochondrial genome (mtDNA) serves as a vital tool in molecular evolution studies, phylogenetic reconstruction and population genetic analyses (Ingman, et al. 2000), which mainly depend on the postulation that mtDNA is strictly maternally inherited. This, in turn, leads to the conclusion that mtDNA is maintained clonally, without the complexity generated from biparental recombination. This assumption, however, has been challenged by investigations that have sparked controversy by providing evidence for paternal inheritance and genetic recombination of mtDNA. The earliest evidence of mtDNA recombination is inferred from population genetic linkage disequilibrium (LD) analysis among pairs of mtDNA polymorphisms (Awadalla, et al. 1999) and phylogenetic analyses of excess homoplasies (homozygous mutations at the same genetic site) for phenotypically silent sites located in mtDNA coding regions (Eyre-Walker, et al. 1999). These analyses showed that LD decreased with distance between alleles and that frequency of homoplasies was much higher than expected on the basis of a single rate of synonymous mutations. Both of these traits are manifestations consistent with recombination. However, these results are not reproducible by using either the same measurement technique (Jorde and Bamshad 2000; Elson, et al. 2001) or an alternative methodology (Jorde and Bamshad 2000). Although reanalysis of mtDNA sequences from the formerly published data sets shows a negative correlation between LD and allele distance, most correlations were not statistically significant or even positive (Piganeau and Eyre-Walker 2004). Similarly, consistent and repeatable results of recombination in a human mtDNA data set cannot be obtained by using six well-established indirect tests of recombination (White and Gemmell 2009). In addition, rather than providing a conclusive argument for the existence of mtDNA recombination (Piganeau and Eyre-Walker 2004), high homoplasmy levels in phylogenetic trees could also be attributed to multiple mutations at hypervariable sites, which may be selected because of their phenotypic advantages. These results reveal a lack of unequivocal proof of mtDNA recombination in the published research using indirect evaluation such as LD analysis and homoplasy evaluation, which may be incapable of determining the very infrequent occurrence of recombination in mtDNA.

Newer, more convincing cases of mtDNA recombination have been documented. Kraytsberg *et al.* reported direct evidence for mtDNA recombination (Kraytsberg, et al. 2004). The authors searched for recombination events in the muscle tissues of an individual with biparental inheritance of the mitochondrial genome (Schwartz and Vissing 2002), which may provide an opportunity for intermolecular recombination between paternal and maternal mtDNA (**Figure 1**). Ultimately, they found 33 recombinants, accounting for ∼0.7% of the total mtDNA molecules. Given the novelty of the result, they went to great lengths to ensure the *in vivo* origin of these recombinants, repeating the experiment with a reconstructed 10:1 mixture of paternal and maternal DNA to exclude the possibility of recombination produced in laboratory (either by PCR or some other errors). According to this result, the frequency of mtDNA recombination is possible but probably extraordinarily low, since the frequency of paternal leakage is rare.

**Figure 1.**
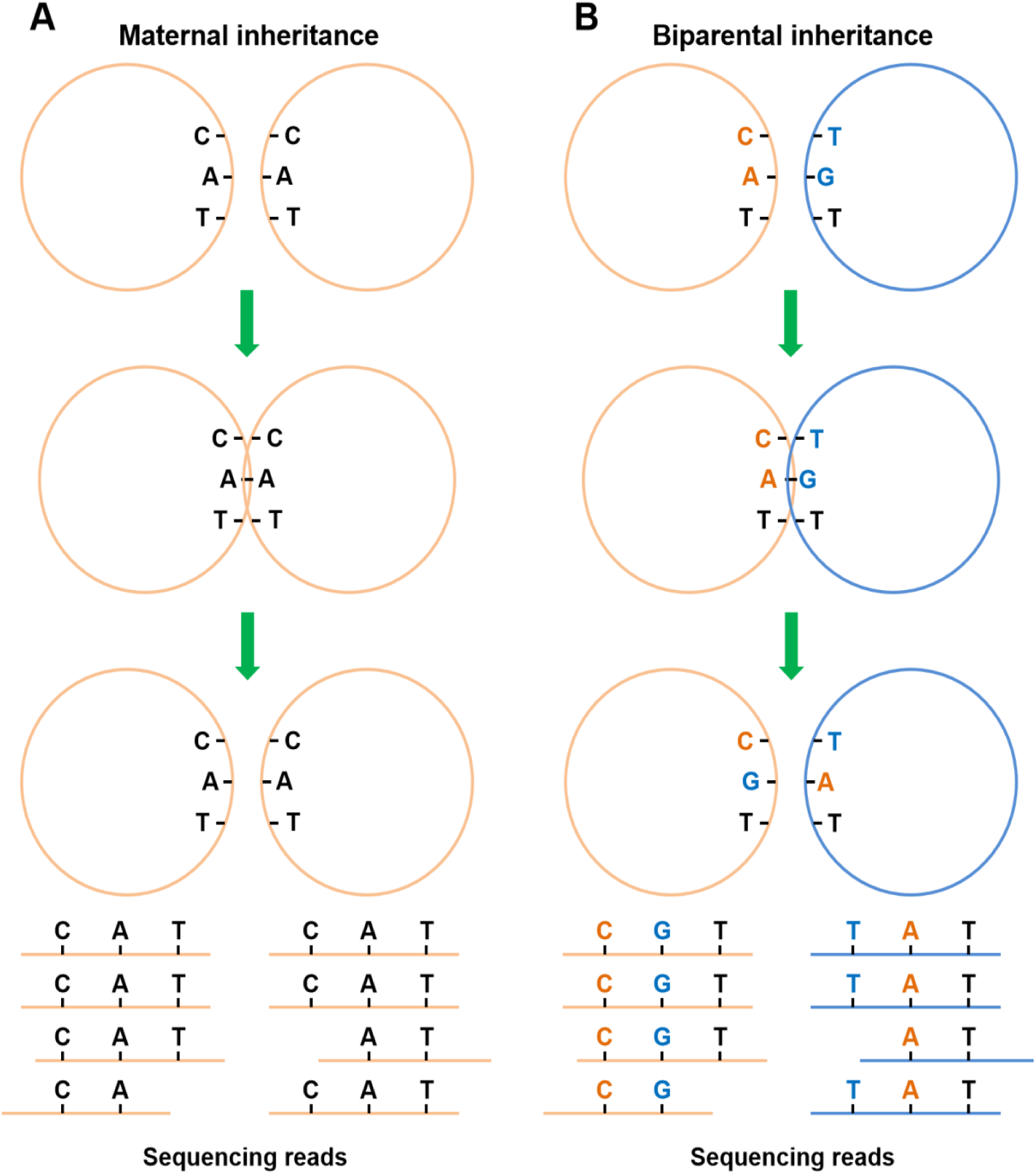
Model of hypothetical recombination between mtDNA molecules. Pink and blue circles represent maternal and paternal mtDNA molecules, respectively. (A) Hypothetical genomic crossover between homoplasmic mtDNA molecules transmitted maternally, which would be expected to rarely generate new haplotypes. (B) Hypothetical recombination between maternal and paternal mtDNA molecules in a biparental inheritance mode. For a detectable recombination event to occur, at least two distinct non-allelic variants must collocate in the mtDNA sequence. In this example, the genotypes C and A are collocated in the maternal mtDNA sequence, while the genotypes T and G are collocated in the paternal sequence. If a crossover event happens (at the locus indicated in the figure), the maternal haplotype turns into C combined with G, while the paternal haplotype changes to T combined with A. After sequencing, different reads covering different regions (as indicated in the figure) are used for genotyping and phasing of the haplotypes.

Further direct evidence of mtDNA recombination between two human cytoplasmic hybrid cell lines supports the idea that recombination is possible in human mtDNA (D’Aurelio, et al. 2004) as evidenced in biparental leakage (Kraytsberg, et al. 2004). The authors fused two human cytoplasmic hybrid cell lines, each containing a distinct pathogenic mtDNA mutation and a specific set of genetic markers used for restriction fragment length polymorphism analysis and relative frequency measurement. Analysis of this hybrid cell model suggested the existence of recombination between the mtDNA haplogroups (∼10% of total mtDNA) (D’Aurelio, et al. 2004). However, these artificially created heteroplasmic cells are in a suboptimal condition for studying recombination, as this condition rarely occurs naturally in humans. Therefore, despite this evidence of recombination, the debate remains.

Individuals with biparental mtDNA transmission will be ideal subjects to test possible intermolecular recombination between paternal and maternal mtDNA. We have identified three unrelated multiple-generation families with a high number and level of mtDNA heteroplasmy (Luo, et al. 2018) after amplifying entire mtDNA by a single long-range PCR and performing next-generation sequencing (NGS) analysis based on an established pipeline (Tang and Huang 2010; Huang 2011; Ma, et al. 2015). For instance, in one of the families (Family A, as shown in **Figure 2**), three immediate relative (lineal relative) members (grandfather-mother-son, or I-1, II-1, and III-2 in the pedigree, respectively) were found to have high levels of mtDNA heteroplasmy (nearly 40% for the minor alleles and 60% for the major alleles). This heteroplasmy covers multiple mtDNA sites in the entire mitochondrial genome (31 sites in both II-1 and III-2, and 20 sites in I-1), which can serve as perfect genetic markers for recombination analysis. These human materials thus provide a unique opportunity to investigate mtDNA recombination.

**Figure 2.**
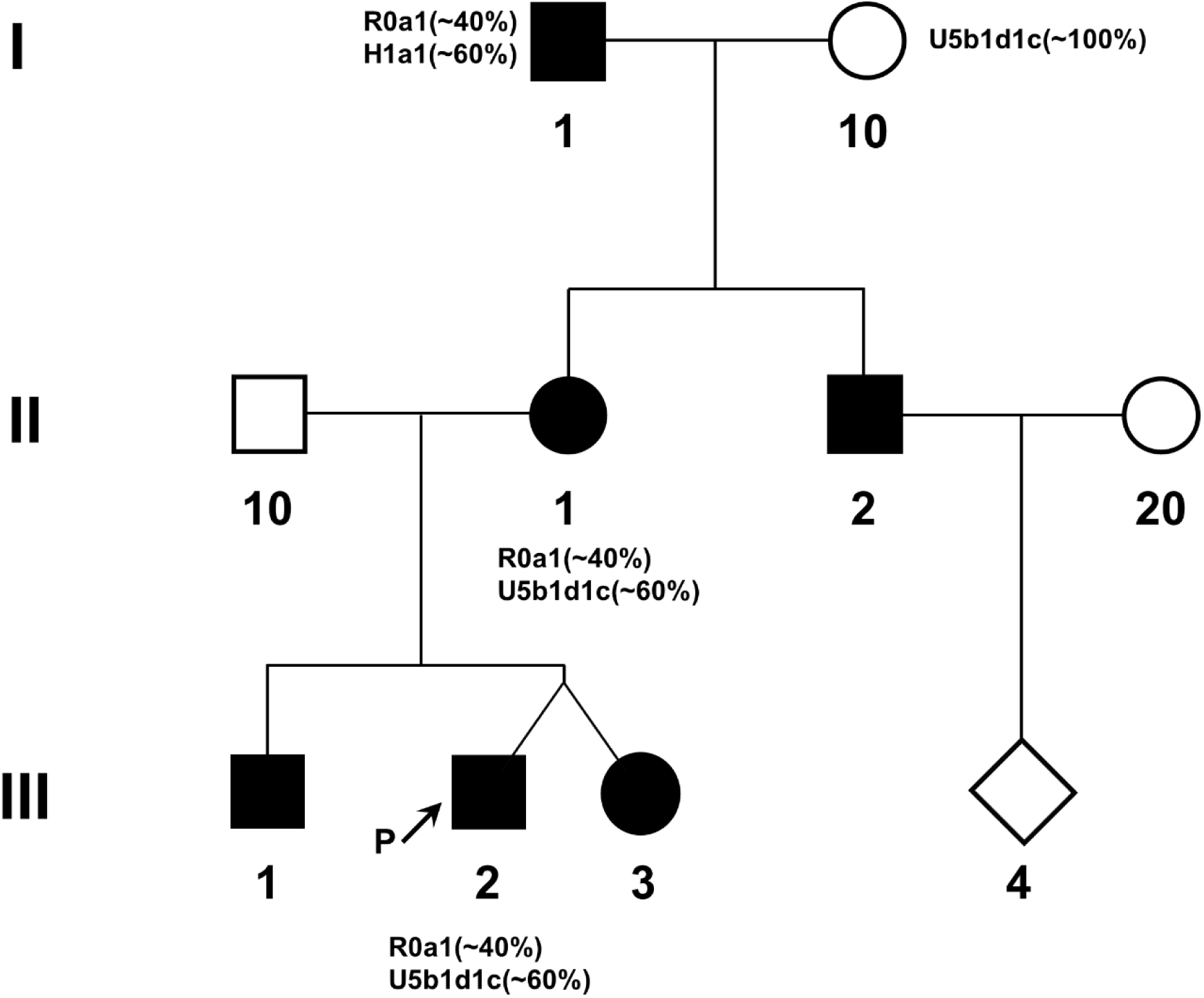
Pedigree of a multiple-generation family (Family A) with a high number and level of mtDNA heteroplasmy. The arrow indicates the proband for the family. Dark squares and circles show individuals with a high level of mtDNA heteroplasmy (nearly 40% for the minor alleles and 60% for the major alleles) at multiple mtDNA sites (31 sites in II-1, III-1, III-2 and III-3, and 20 sites in I-1). Their haplogroups and the corresponding average heteroplasmy ratio are indicated. For example, the haplogroup R0a1 and H1a1 account for ∼40% and ∼60% of the mtDNA, respectively, in I-1. For SMRT sequencing, whole mitochondrial DNA was PCR-amplified from the blood samples of I-1, II-2, and III-2 (**Table 1**), as well as from the fibroblasts of I-1 (**Figures 7-8**).

In this study, we first investigated the possibility of mtDNA recombination by sequencing the entire mtDNA from blood cells of three immediate relative individuals (I-1, II-1, and III-2) with biparental mtDNA transmission in Family A, using single-molecule real-time (SMRT) sequencing technology. The SMRT sequencing technology allows us to revisit the controversial issue of mtDNA recombination accurately by determining the phase of all observed single nucleotide polymorphisms (SNPs) across all or nearly all of the entire 16.5-kbp mitochondrial genome at a single-molecule level, without cross-referencing between different molecules (as occurs in Sanger or NGS). After phasing the genotypes of mtDNA SNPs, no evidence of intermolecular recombination between paternal and maternal mtDNA was found, as shown in **Table 1**. For the immediate relative grandfather (I-1), two distinct haplogroups, R0a1 and H1a1, were identified after phasing of the 17 called genotypes with an average genotype frequency of ∼40% and ∼60%, respectively. The haplogroup R0a1 was transmitted to his daughter (II-1), who inherited another haplogroup, U5b1d1c, from her mother (I-10). No evidence of recombination between the R0a1 and H1a1 haplogroup was found in the grandfather (I-1) when transmitting the mtDNA in a biparental pattern. Similarly, when maternally transmitting the biparental mtDNA from the mother (II-1) to the son (III-2), no recombination appears to have occurred.

**Table 1.**
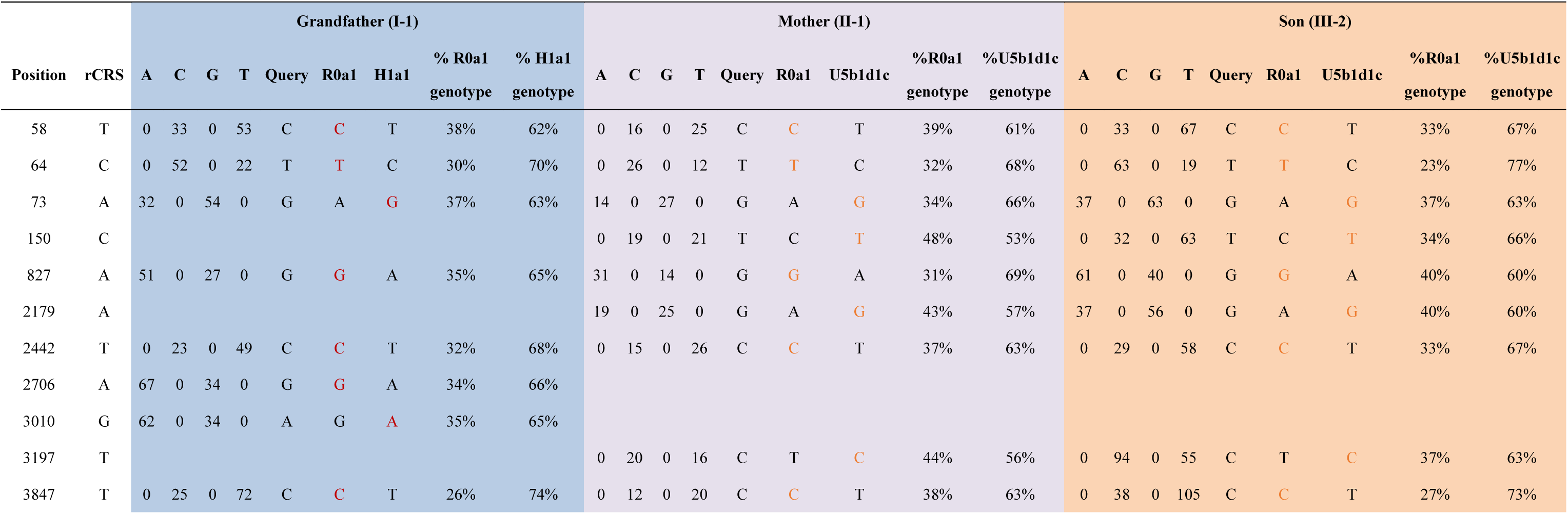
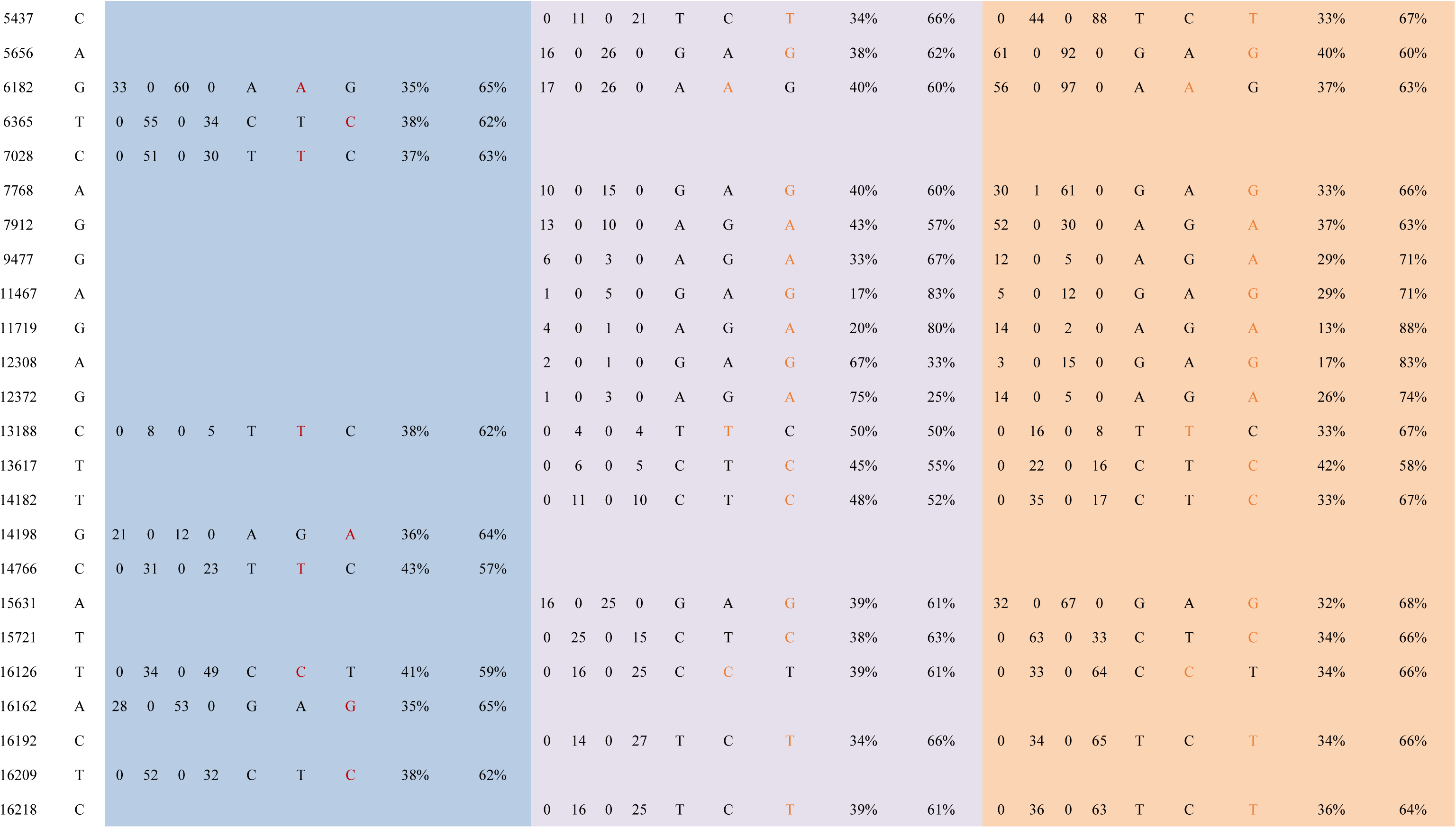
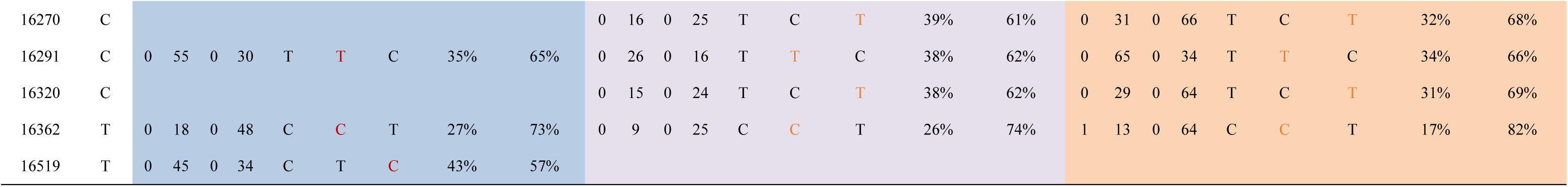
Phasing of the haplogroups in the three immediate relative individuals from Family A. The query genotypes found in specific individuals represent genotypes different from the reference sequence (rCRS, or “Revised Cambridge Reference Sequence”). The rCRS is the corrected human mtDNA reference sequence, which is deposited in the GenBank NCBI database under accession number NC_012920. The percentage of specific genotypes for different mtDNA markers varied mainly because they are covered by different sequencing reads, resulting in different read depth for specific loci.

However, this initial analysis was lacking in certain regards. First, this initial dataset was generated when these families were first identified, and thus the data were generated using older PacBio technology with a much higher per base error rate and shorter average reads. The longest single molecules from this dataset were about 12 kbp, and the average read was around 5 kbp. There were relatively few of the ∼12 kbp reads, and none of the individual single molecular reads fully covered the entire 16.5 kbp mitochondrial genome. These incomplete full mtDNA reads, while certainly better than the several hundred bases provided by traditional NGS sequencing methods, make this data less than ideal for detecting recombination, since there are much fewer variant sites on each continuous sequence. The high error rate and lower coverage also decrease the accuracy of base calls, and recombination hotspot analysis is likely to be heavily skewed due to very uneven coverage in regions far from the primer start site. There is also a more conceptual question related to the possibility of tissue-specific mtDNA recombination differences. It is possible that mtDNA recombination only occurs in particular cell lineages, and that some lineages experience high rates of mtDNA recombination while others experience no mtDNA recombination at all. Thus, it is important that we test DNA derived from tissues other than blood in order to maximize our chance of detecting mtDNA recombination events.

For these reasons, we obtained three new DNA samples for a new round of SMRT sequencing analysis based on new chemistry and technology (see Materials and Methods). The fibroblast DNA sample was included from individual I-1 of Family A. The other two samples were blood DNA samples obtained from individuals belonging to two other families showing biparental mtDNA transmission: Family B (Individual I-1, **Figure 3**) and Family C (Individual III-1, **Figure 4**). As before, all three DNA samples were subjected to long-range PCR amplification of their full-length mtDNA, and the resulting amplicons were submitted for SMRT sequencing. The resulting reads demonstrated this new approach has a much better average read length and accuracy. Most importantly, for each sample, we were able to obtain a minimum of 1000 full-length reads that span the full length (16,569 bp) of the mitochondrial genome (1712 reads for the fibroblast sample from Family A I-1, 2160 reads for Family B I-1, and 2719 reads for Family C III-1) (**Table 2**). This allowed for a much more refined analysis of recombination hotspots and recombination frequency.

**Table 2.** Number of parental and recombinant sequences for each sample. A custom script was used to look at all of the variant positions for each sample and phase sequences into their mtDNA haplogroups. For this table, a “breakpoint” is defined as a switch from one parental haplogroup to the other that is sustained for at least two sites.

|  | **Family A, I-1** |  | **Family B, I-1** |  | **Family C, III-1** |  |
| --- | --- | --- | --- | --- | --- | --- |
|  | # of reads | Percent % | # of reads | Percent % | # of reads | Percent % |
| Total | 1712 | 100 | 2160 | 100 | 2719 | 100 |
| Paternal | 12 | 0.70093458 | 75 | 3.47222222 | 1130 | 41.5593968 |
| Maternal | 1234 | 72.0794393 | 908 | 42.037037 | 629 | 23.133505 |
| 0 Breakpoints | 1246 | 72.7803738 | 983 | 45.5092593 | 1759 | 64.6929018 |
| 1 Breakpoint | 141 | 8.23598131 | 406 | 18.7962963 | 488 | 17.9477749 |
| 2 Breakpoints | 208 | 12.1495327 | 278 | 12.8703704 | 291 | 10.7024641 |
| 3 Breakpoints | 63 | 3.67990654 | 199 | 9.21296296 | 126 | 4.63405664 |
| 4 Breakpoints | 43 | 2.51168224 | 143 | 6.62037037 | 41 | 1.50790732 |
| 5 Breakpoints | 9 | 0.52570093 | 81 | 3.75 | 13 | 0.47811695 |
| 6 Breakpoints | 2 | 0.11682243 | 42 | 1.94444444 | 1 | 0.03677823 |
| 7 Breakpoints |  |  | 17 | 0.78703704 |  |  |
| 8 Breakpoints |  |  | 7 | 0.32407407 |  |  |
| 9 Breakpoints |  |  | 4 | 0.18518519 |  |  |

**Figure 3.**
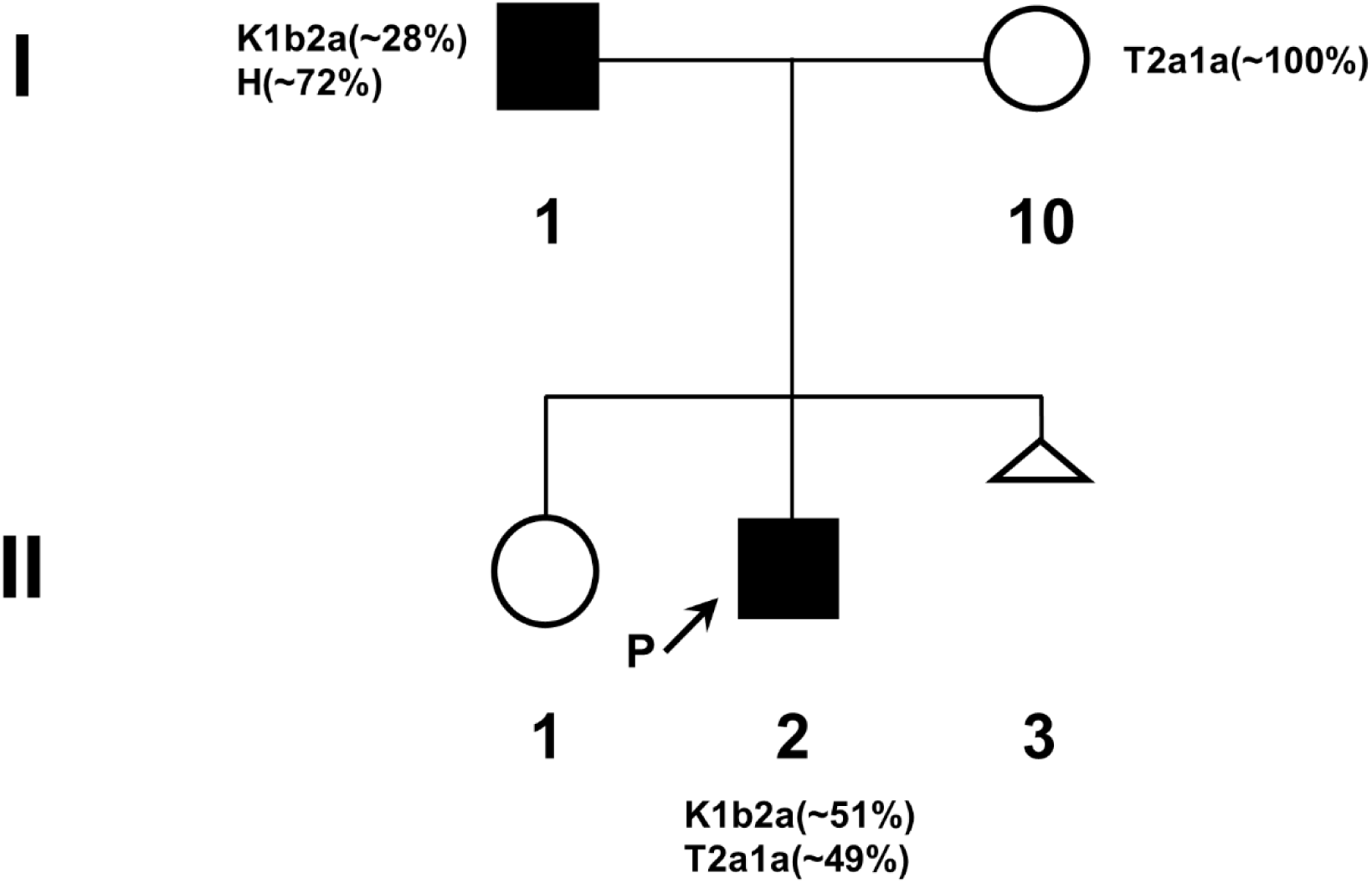
Pedigree of a multiple-generation family (Family B) with a high number and level of mtDNA heteroplasmy. The arrow indicates the proband for Family B. Dark squares and circles show individuals with a high level of mtDNA heteroplasmy at multiple mtDNA sites (29 sites in I-I and 44 sites in II-2). Their haplogroups and the corresponding average heteroplasmy ratio are indicated. For SMRT sequencing, whole mitochondrial DNA was PCR-amplified from the blood sample of I-1, the proband’s father (**Figures 7-8**).

**Figure 4.**
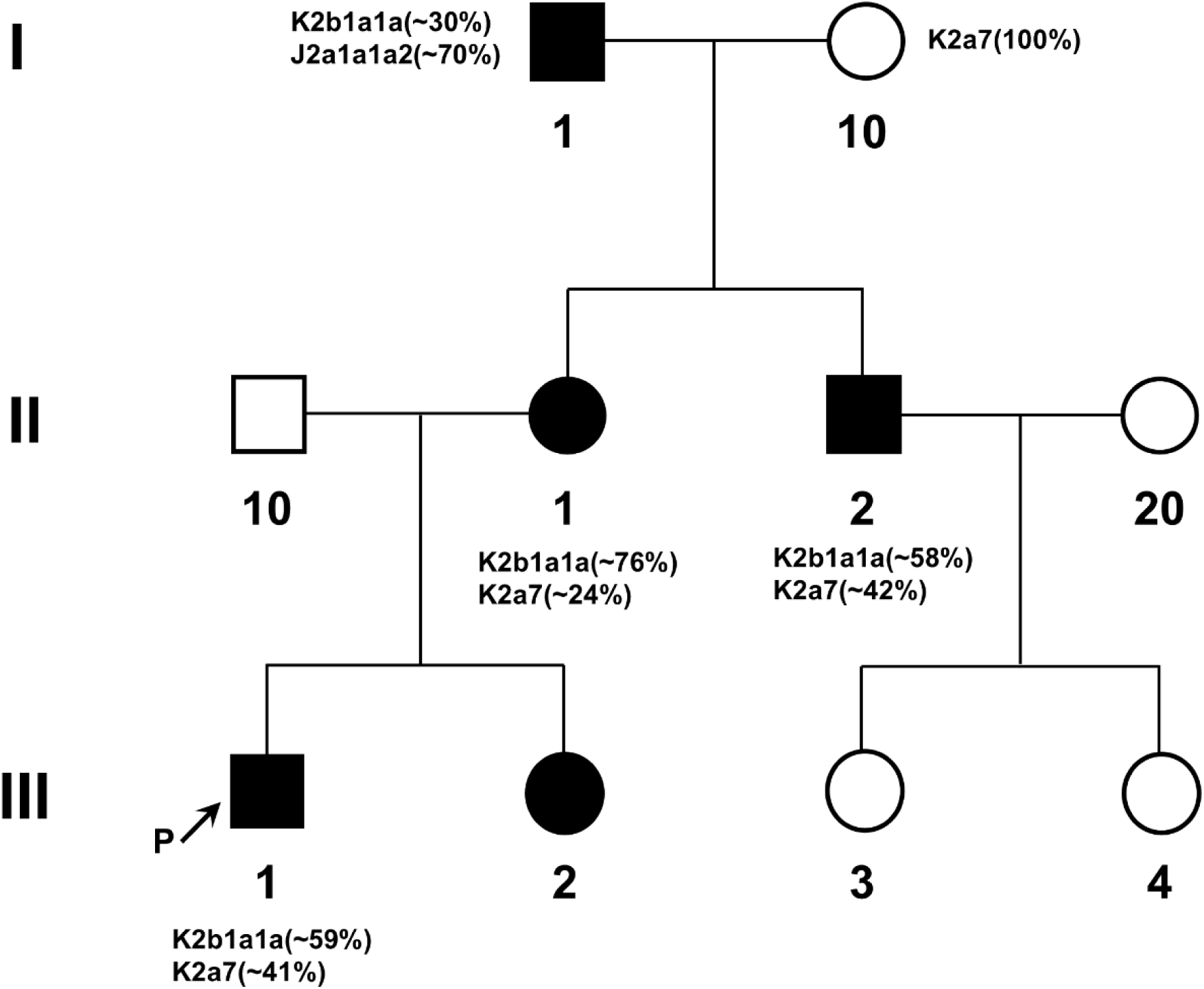
Pedigree of a multiple-generation family (Family C) with a high number and level of mtDNA heteroplasmy. The arrow indicates the proband for Family C. Dark squares and circles show individuals with a high level of mtDNA heteroplasmy at multiple mtDNA sites (51 sites in I-1 and 13 sites in II-1, II-2, III-1, and III-2). Their haplogroups and the corresponding average heteroplasmy ratio are indicated. For SMRT sequencing, whole mitochondrial DNA was PCR-amplified from the blood sample of III-1, the proband (**Figures 5-8**).

As expected, our results demonstrated the existence of two full-length mtDNA species corresponding to the predicted maternal and paternal haplogroups for each of the samples (**Figure 5**), further confirming the conclusions of our original study of biparental mtDNA transmission (Luo, et al. 2018). Curiously, our results also showed that a large proportion of reads contained at least one putative crossover event between the maternal and paternal haplotypes (**Figure 6**). However, a deeper analysis indicates that these recombination events are very likely spurious, for the following reasons. First, the distribution of breakpoints (i.e. putative “crossover” sites) in recombinants for all three samples was near linear, with no apparent recombination hotspots (**Figure 7**). This result would seem to suggest that recombination is random, which conflicts with literature showing breakpoint hotspots at sites with biological significance (Kraytsberg, et al. 2004). The number of recombinants found in our samples, 28-55% of total mtDNA, is also much higher than the rates of 0.7% in Kraytsberg et al. A likely explanation is that the recombinants were generated by PCR and not *in vivo* (Lahr and Katz 2009). Modeling of PCR chimera formation rates shows a similar distribution of recombinants and non-recombinants as observed in our actual data (**Figure 8**), further suggesting that these events are PCR artifacts rather than true, *in vivo* recombination events. It should also be noted that the ratio of purely paternal to purely maternal mtDNA sequences was different from the heteroplasmy levels obtained from short-read sequencing (**Table 2**). In the phased PacBio CCS data, the majority parental haplogroup (whether paternal or maternal) accounted for a greater portion of the full-length non-recombinant sequences than expected from previous variant calling. A possible explanation for this is that PCR-based recombination would shift the proportion of unrecombined reads in favor of the majority haplogroup. In the case where maternal sequences outnumber paternal sequences, both sequences have the same chance of failing to fully extend (the most common cause of PCR-based recombination). However, a partially extended sequence is more likely to anneal to a maternal template, since in this case, maternal sequences are more numerous. Under these conditions, when a maternal sequence undergoes PCR-based recombination, it is very likely to recombine to another maternal template, which would generate a purely maternal sequence. When the paternal sequence undergoes PCR-based recombination, however, it is also likely to recombine to a maternal sequence, generating a recombinant sequence. This shift can be seen in both cases where the maternal sequence is the majority haplogroup (Family A and B) and in the case where the paternal sequence is the majority (Family C). Thus, on balance, the evidence obtained from our SMRT sequencing data indicates that these “recombination” events are PCR artifacts, and that true *in vivo* recombination is exceedingly rare, if not entirely non-existent.

**Figure 5.**
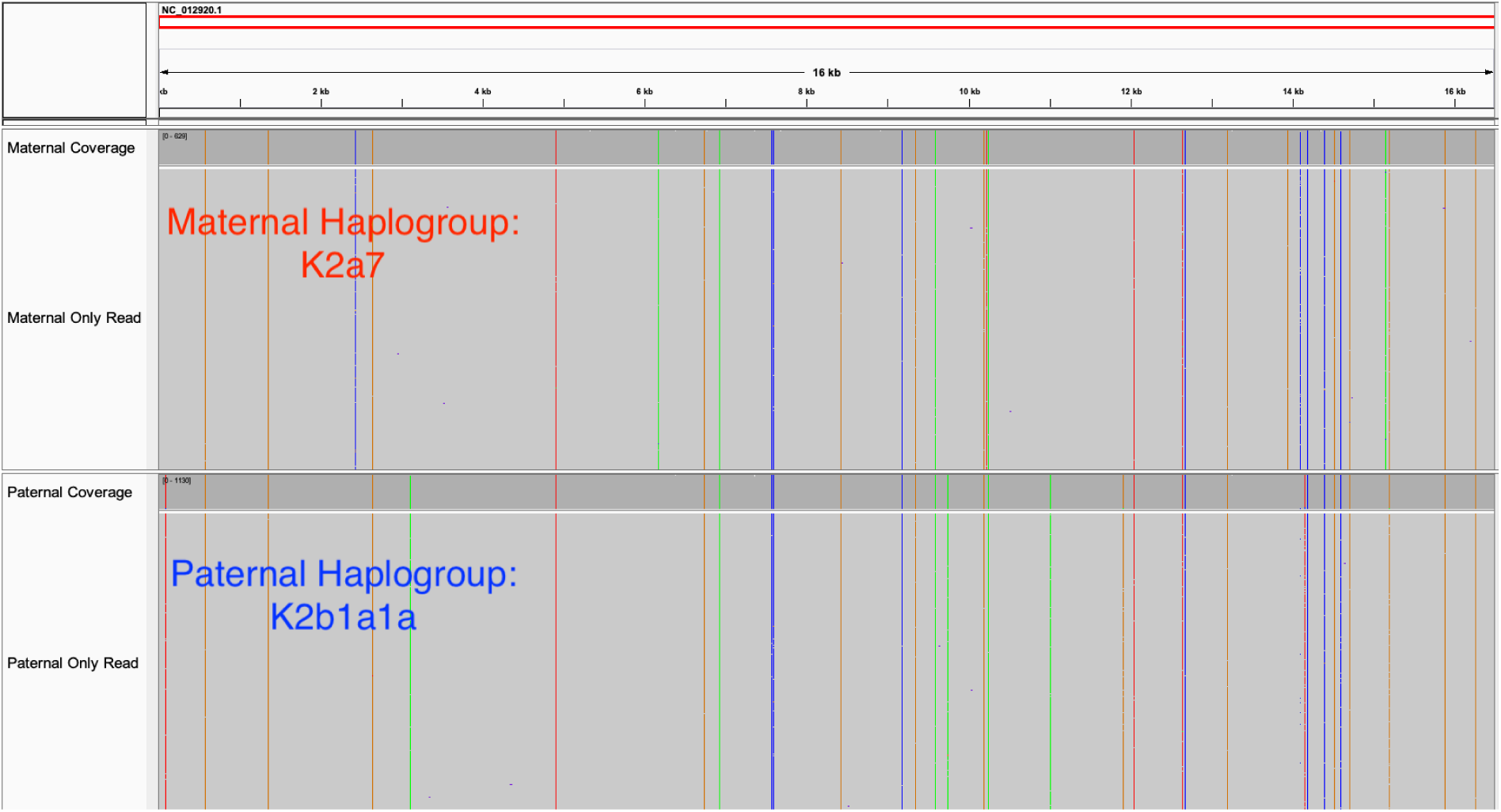
Comparison of sequence reads for parental haplogroups of III-1 of Family C. This diagram shows the non-recombinant reads from the original parental haplogroups for III-1 from Family C. The vertical bars with different colors represent different base substitutions when compared to the rCRS reference genome (NC_012920). The maternal haplogroup is K2a7, and the paternal haplogroup is K2b1a1a. The CCS reads are aligned to the modified reference file, filtered for quality and length, then visualized in Integrative Genomics Viewer (IGV) software.

**Figure 6.**
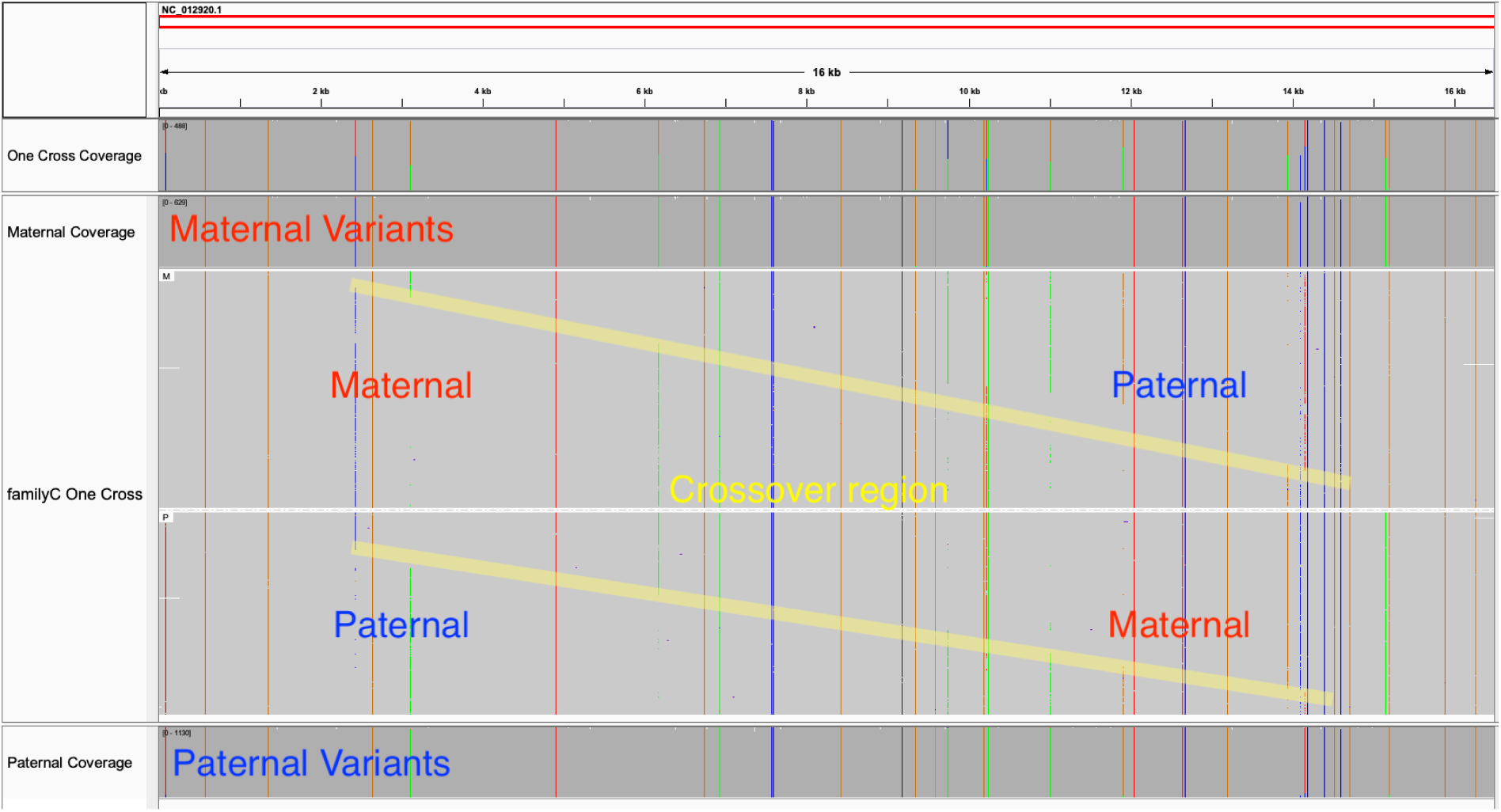
Recombinant sequences with one breakpoint from III-1 in Family C. This diagram shows all of the single-breakpoint reads from III-1 in Family C. The yellow band indicates the location where the sequences switch from one haplogroup to the other. The vertical bars with different colors represent different base substitutions when compared to the rCRS reference genome (NC_012920). For comparison, non-recombinant reads for the maternal haplogroup are shown at the top of the diagram, and non-recombinant reads for the paternal haplogroup are shown at the bottom. As in **Figure 5**, the original maternal haplogroup is K2a7, and the paternal haplogroup is K2b1a1a.

**Figure 7.**
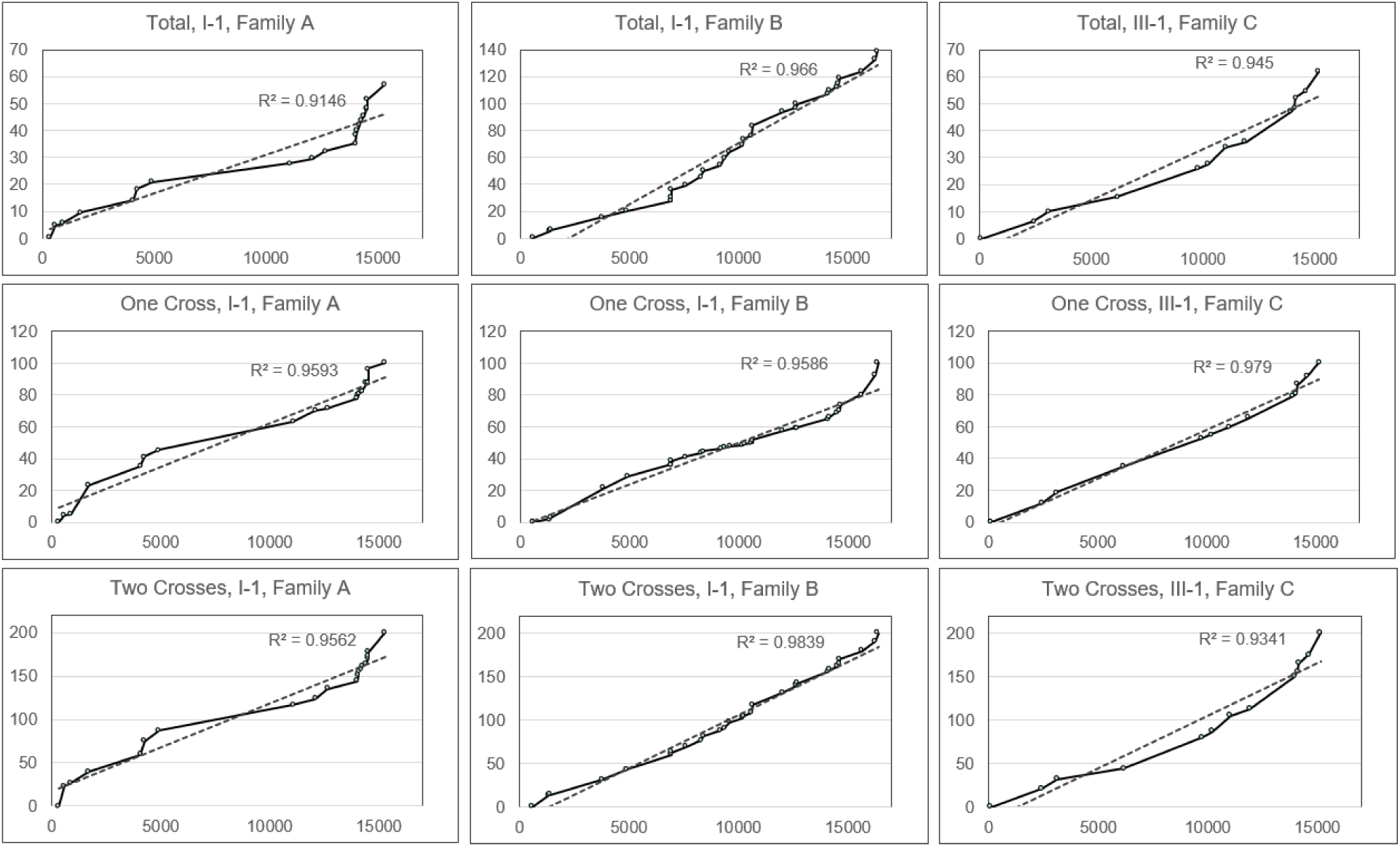
Map of breakpoint sites and frequency. These graphs show the cumulative percentage of reads with a breakpoint at each site in the mitochondrial genome. A custom script was used to count the number of strands with a breakpoint at each variant site. The rates are cumulative to normalize for the distance between variant sites. Position “0” below corresponds to position “2120” in the rCRS, which is the annealing site of the forward primary used to generate the PCR amplicon used for sequencing. In the same way, position “16569” below corresponds to “2119’ in the rCRS, which is the annealing site of the reverse primer used to generate the amplicon.

**Figure 8.**
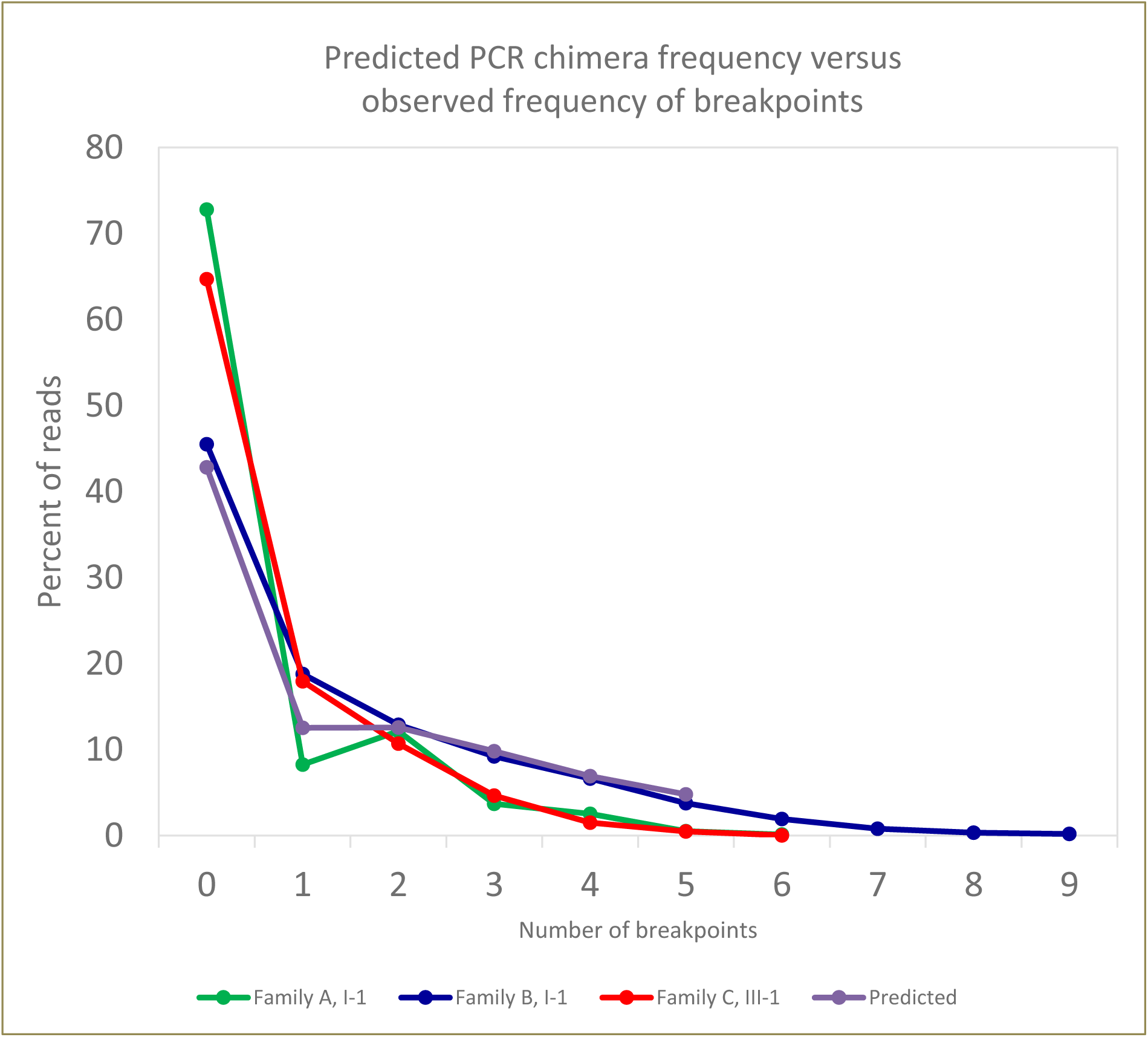
Modeling of predicted PCR chimera rates relative to observed breakpoint frequencies. This graph shows the distribution of reads according to the number of breakpoints observed in the SMRT sequencing data for the fibroblasts from I-1 of Family A, as well as the blood samples from I-1 of Family B and III-1 of Family C. The predicted distribution for PCR-generated chimeras is plotted for comparison.

By performing SMRT sequencing across multiple families and tissue types, our study further indicates the absence or lack of intermolecular mtDNA recombination in humans when the mtDNA is transmitted to offspring in either biparental or maternal mode. Our evidence comes from direct observation of recombination events in lineal relative members, rather than indirect population genetic inference. Furthermore, multiple informative SNPs from around the entire mtDNA were called for genotyping and phasing of the haplogroup, eliminating errors due to phasing mistakes and providing solid support for the argument. For a detectable mtDNA recombination event, it is necessary that at least two distinct non-allelic variants colocate in the same mtDNA sequence (**Figure 1**). Human mitochondrial genomes possess low-frequency rearrangements, i.e., partial duplication or deletion, generated by intramolecular recombination on break-point hotspot regions often defined by short direct repeats (Kajander, et al. 2000). However, little evidence has been found indicating the possibility of intermolecular recombination in human mtDNA. A plausible manner that mtDNA recombination could influence genetic diversity is by exchanging genetic information between biparental molecules at the stage of fertilization and pre-implantation embryo development, during which the paternal mtDNA may be eliminated gradually by active mechanisms of selective ubiquitin-dependent protein degradation and/or mitophagy (Song, et al. 2016). Even if paternal mtDNA fails to be effectively eliminated, resulting in biparental heteroplasmy (Schwartz and Vissing 2002; Luo, et al. 2018) and providing the potential for genomic exchanges, the occurrence of recombination between the paternal mtDNA and maternal mtDNA requires physical proximity of the two mitochondrial genomes. However, mammalian mitochondria typically only contain a single copy of mtDNA (Kukat, et al. 2011), which reduces the chance of recombination events between two molecules expected to be within close proximity. Additionally, no definitive evidence has been discovered to indicate whether or not paternal and maternal heterologous mitochondria can fuse together to facilitate the proximity of the biparental mtDNA and exchange of their genetic material.

Intermolecular mtDNA recombination has been reported in some taxa, but is still controversial in others. There is increasing evidence confirming that mtDNA undergo recombination in plants (Stadler and Delph 2002; Bergthorsson, et al. 2003; Apitz, et al. 2013) and fungi (Saville, et al. 1998), and such events readily occur even in yeast (MacAlpine, et al. 1998; Fritsch, et al. 2014). However, it is still difficult to come to a definite conclusion for widespread animal mtDNA recombination (Greiner, et al. 2015). Direct evidence for recombination of mtDNA in species like the nematode *Meloidogyne javanica* (Lunt and Hyman 1997), the mussel *Mytilus galloprovincialis* (Ladoukakis and Zouros 2001a), and the flatfish *Platichthys flesus* (Hoarau, et al. 2002) has been well documented. It is worth noting that intermolecular recombination should be much easier to observe in species which do not follow strict uniparental inheritance of mitochondria. For example, the mussel *Mytilus galloprovincialis* possesses two lineages of mtDNA, each with a distinct transmission mode: a phenomenon known as doubly uniparental inheritance (DUI) (Zouros, Ball, et al. 1994; Zouros, Oberhauser Ball, et al. 1994). Several computational statistical analyses comparing the published animal mtDNA sequences also provided evidence for recombination even in organisms with standard maternal mtDNA inheritance (Ladoukakis and Zouros 2001b; Tsaousis, et al. 2005). However, when using different methods to test the potential recombination events of these mtDNA sequences, no or only weak evidence could be found in many of these species (Smith and Smith 2002). A recent study used an *in vivo* mouse model with germline heteroplasmy for a defined set of mtDNA mutations for more than 50 generations to detect potential recombination events (Hagstrom, et al. 2014). Based on cloning of single mtDNA molecules in the λ phage (without prior PCR amplification), followed by subsequent mutation analysis, no germline recombination was found after transmission of mtDNA under genetically and evolutionarily relevant conditions. These findings provide similar evidence as what was observed in our study; that is, the distinct populations of mtDNA carried by these mice did not show any observable evidence of recombination when transmitted maternally.

In summary, the results in our study largely deny the occurrence of intermolecular mtDNA recombination in both biparental and maternal transmission modes, thus supporting the feasibility of mtDNA-based molecular evolution studies, phylogenetic inference and population genetic analyses for humans, while also avoiding spurious phylogenetic inferences and incorrect rejection of the molecular clock. Furthermore, these findings also broaden our knowledge of human mtDNA-associated diseases. However, the question of intermolecular mtDNA recombination remains far from resolved, and more evidence and critical discussion are warranted to come to a definite conclusion and to develop our understanding of human and mitochondrial evolution.

## Material and Methods

### Subjects

Three unrelated multi-generation families with a high number and level of mtDNA heteroplasmy were previously identified (Luo, et al. 2018). For this study, heteroplasmic individuals from each of these families were chosen for further analysis. For Family A, three immediate related individuals (lineal relatives) across multiple generations were chosen: a grandfather (I-1), mother (II-1), and son (III-2), as shown in the pedigree in **Figure 2**. For Family B and C, a single individual was chosen from each family for analysis: I-1 for Family B (**Figure 3**), and III-1 for Family C (**Figure 4**). Family A and Family B were originally evaluated at the MitoClinic and referred to the Mitochondrial Diagnostic Laboratory at the Human Genetics Division of Cincinnati Children’s Hospital Medical Center (CCHMC), and Family C was evaluated at Mayo Clinic and genetic testing was done at the Diagnostic Laboratory at Baylor College of Medicine. For all families, mtDNA analysis was performed based on long-range PCR and next-generation sequencing (Tang and Huang 2010; Huang 2011; Ma, et al. 2015; Luo, et al. 2018). To investigate the possibility of recombination events in these individuals’ mtDNA, written informed consent was obtained from all participants. This study was approved by the Institutional Review Board of CCHMC (approval study ID: 2013-7868).

### Whole mitochondrial DNA amplification

Genomic DNA was isolated from the blood and fibroblast samples using Gentra DNA extraction kit (Qiagen, Hilden, Germany) according to the manufacturer’s instructions. Entire mtDNA was amplified as a single, long-range PCR amplicon as previously described (Tang and Huang 2010; Huang 2011; Ma, et al. 2015). One hundred nanograms of total genomic DNA were used as template in a 50-μL PCR system. The primers specifically recognize genuine mtDNA: F-2120 (GGACACTAGGAAAAAACCTTGTAGAGAGAG) and R-2119 (AAAGAGCTGTTCCTCTTTGGACTAACA). PCR amplifications were performed using TaKaRa LA Taq Hot Start polymerase (TaKaRa Biotechnology, Kyoto, Japan). PCR conditions were: 94 °C for 1 min; 98 °C for 10 sec and 68 °C for 16 min, 30 cycles; 72 °C for 10 min and hold at 4 °C.

### Single-molecule real-time (SMRT) sequencing and analysis

The long-range PCR products of entire mtDNA molecules were used for SMRT sequencing at the Institute of Genome Science, University of Maryland for the blood DNA samples for I-I, II-1, and III-2 of Family A, and at Novogene (Beijing, China) for the fibroblast DNA from I-1 of Family A and the blood DNA for I-1 in Family B and III-1 in Family C.

For the data generated at the University of Maryland, DNA libraries were constructed and prepared for sequencing using the SMRTbell Template Prep Kit 2.0 and the DNA/Polymerase Binding Kit 2.0 (Pacific Biosciences, Menlo Park, CA) according to the manufacturer’s instructions. 16.5-kbp fragments were size-selected with the BluePippin device (Sage Sciences, Beverly, MA), and were sequenced with the DNA Sequencing Kit 2.0 (Pacific Biosciences, Menlo Park, CA) using a 4-hour movie collection. The raw subread data was analyzed using PacBio’s Long Amplicon Analysis module (Smrtanalysis 2.3.0) to find phased consensus sequences, using PacBio subreads that were longer than 13 kbp. The resulting phased sequences were aligned to the mitochondrial reference genome and variant calling was performed for each. The variants were then used to assign a genotype to each phased sequence by comparing them with variants found in the parental/ancestral mitochondrial genomes (Kurtz, et al. 2004). Circular consensus sequence (CCS) were also generated from the amplicon data and used to calculate allele/haplogroup frequencies (Chaisson and Tesler 2012).

For the data generated at Novogene, DNA libraries were constructed and prepared for sequencing using the SMRTbell Template Prep Kit 2.0 and the Sequel II Binding Kit 2.0 (Pacific Biosciences, Menlo Park, CA) according to the manufacturer’s instructions. The PCR amplicon were sequenced without size-selection using PacBio’s Sequel System (Pacific Biosciences, Menlo Park, CA), using Chemistry 3.0, Software version 6.0, and a 10-hour movie collection time. The raw subread data was used to generate consensus sequences using pbccs, a command-line tool provided by PacBio. All three data sets were run with minimum subread length at 13 kbp, filtering out shorter reads. Due to the position of the primers, full-length reads started at base position 2120 of the reference NC_012920 genome and ended at 2119. To facilitate accurate alignment of these long reads, the reference mtDNA sequence was modified by moving bases 1-2119 to the end of the reference sequence. The consensus reads were then aligned to the modified sequence using pbalign, also part of the PacBio command-line tools. Using a quality filter of RQ >= .995 and minLength = 16000, the aligned consensus reads were reduced to only high-quality reads that are full-length or nearly full-length. A custom script was then used to separate the aligned reads into parental haplogroups and recombinant groups. We used the SNP sites that differed between the two parental haplogroups to count the number of paternal and maternal variants on each read. Using the pattern of variant haplotypes, we sorted the reads into their respective groups. The position and haplotype of the sites were based on the original analysis of the short-read data for these families (Luo, et al. 2018). Each CCS sequence was assigned to either the paternal or maternal haplogroup based on the first heteroplasmic variant site. Each site where the paternal and maternal haplogroups disagreed was checked to see if it was paternal or maternal. If variants switched haplogroup and the new haplogroup was sustained for at least two variant sites, then the CCS sequence was marked as recombinant and a crossover at the site recorded. The reads were visualized and checked in Integrative Genomics Viewer (IGV) software (Robinson, et al. 2011). A map of the breakpoint sites was generated to check for potential recombination hotspots.

The model of PCR recombinant distribution was generated based on the assumption that for each round of PCR, a certain percentage of sequences would not fully extend. The template to which the partially extended DNA strand anneals is determined by the distribution of sequences after the previous round of PCR. The final distribution of parental and recombinant sequences after 30 rounds of PCR was compared to the observed distribution of sequences in the three samples. A predicted distribution was generated for all combinations of heteroplasmy level and recombination rates, and the results used to generate chi-square statistics to find the combination of best fit.

For all datasets, the mitochondrial haplogroups were identified based on defining SNP genotypes provided by SNPedia (https://www.snpedia.com/index.php/MtDNA_Haplogroup) and MITOMAP (http://www.mitomap.org/foswiki/bin/view/MITOMAP/HaplogroupMarkers).

## Acknowledgements and funding

The authors would like to thank the subjects for participating in our study. We also thank the Institute of Genome Science, University of Maryland, Novogene, and the DNA Core facility of CCHMC, where the SMRT sequencing and next-generation sequencing were performed. This work was supported by the Cincinnati Children’s Hospital Research Foundation and the Eunice Kennedy Shriver National Institute of Child Health & Human Development at the National Institute of Health (1R01HD092989-01A1) to T.H.

